# Age, sex, and social environmental effects on immune cell composition in a free-ranging non-human primate

**DOI:** 10.1101/2021.12.06.471383

**Authors:** Mitchell R. Sanchez Rosado, Nicole Marzan-Rivera, Marina M. Watowich, Andrea D. Negron-Del Valle, Petraleigh Pantoja, Melissa A. Pavez-Fox, Erin R. Siracusa, Eve B. Cooper, Josue E. Negron-Del Valle, Daniel Phillips, Angelina Ruiz-Lambides, Cayo Biobank Research Unit, Melween I. Martinez, Michael J. Montague, Michael L. Platt, James P. Higham, Lauren J. N. Brent, Carlos A. Sariol, Noah Snyder-Mackler

## Abstract

Increasing age is associated with dysregulated immune function and increased inflammation– patterns that are also observed in individuals exposed to chronic social adversity. Yet we still know little about how social adversity impacts the immune system and how it might promote age-related diseases. Here, we investigated how immune cell diversity varied with age, sex and social adversity (operationalized as low social status) in free-ranging rhesus macaques. We found age-related signatures of immunosenescence, including lower proportions of CD20+ B cells, CD20+/CD3+ ratio, and CD4+/CD8+ T cell ratio – all signs of diminished antibody production. Age was associated with higher proportions of CD3+/CD8+ Cytotoxic T cells, CD16+/CD3-Natural Killer cells, CD3+/CD4+/CD25+ and CD3+/CD8+/CD25+ T regulatory cells, and CD14+/CD16+/HLA-DR+ intermediate monocytes, and lower levels of CD14+/CD16-/HLA-DR+ classical monocytes, indicating greater amounts of inflammation and immune dysregulation. We also found an effect of exposure to social adversity (i.e., low social status) that was sex-dependent. High-status males, relative to females, had higher CD20+/CD3+ ratios and CD16+/CD3 Natural Killer cell proportions, and lower proportions of CD8+ Cytotoxic T cells. Further, low status females had higher proportions of cytotoxic T cells than high status females, while the opposite was observed in males. High status males had higher CD20+/CD3+ ratios than low status males. Together, our study identifies immune cell types that differ by age in a human-relevant primate model animal, and demonstrates a novel link between sex-dependent immunity and social adversity.

## Introduction

The average human lifespan has almost doubled over the past century [1], accompanied by an increase in the prevalence of many age-related diseases, including cardiovascular disease, autoimmune disease, diabetes, arthritis, and cognitive decline [2–5]. As individuals age, there is a disruption in the homeostatic balance between innate and adaptive immunity linked to both increases in age-related disease and susceptibility to infection. This imbalance is reflective of two age-related alterations, namely increased inflammation (“inflammaging”) and a decline in adaptive immune function (“immunosenescence”) [6–7]. Both alterations disrupt the balance between pro- and anti-inflammatory mediators that characterize a healthy immune system. For example, with increasing age, innate immune cells such as monocytes become more active and release drivers of inflammation that include proinflammatory cytokines (e.g., TNF-α) [8]. Adaptive immune cells, such as B cells and helper T cells, show age-related declines that directly impact long-term immunity, as exemplified by the lower effectiveness of vaccination in older individuals [9].

Yet these age-related alterations in immunity are not universal in their trajectories across individuals. There is substantial heterogeneity with age; not all individuals age at the same rate or fall victim to the same age-related diseases. For instance, some people become hypertensive in their 30s, while a 60-year-old may never suffer from this condition. Part of this heterogeneity is due to sex differences in the immune system, which alter the prevalence and onset of age-related diseases. For example, in many species, females mount a stronger immune response with increasing age compared to males [10–11]. Females also have a stronger age-related increase in inflammatory cells compared to males [12]. Further, in humans, older men are more susceptible to infections, such as leptospirosis, tuberculosis, and hepatitis A, than are women [13].

Heterogeneity in aging can also arise from life experiences, such as exposure to social adversity, which can influence the onset and progression of disease and, ultimately, mortality [14]. Social adversity, which is often associated with low social status, and other social stressors [15–16], has been linked to accelerated aging as indexed by biomarkers like epigenetic age and telomere attrition [17–19]. There are also broad similarities between the effects of age and social adversity on peripheral immune function [20]. For instance, early life social adversity in humans has been linked to increases in proinflammatory T cells [21] – a characteristic usually seen with increasing age. Further, various adversities and social stressors are associated with a decrease in naïve CD4 T helper cells and an increase in naïve CD8 T cells [22], pointing to how the social environment can shape immunity. However, the extent to which social adversity may be associated with immunity across the lifecourse remains unknown. Social adversity might lead to accelerated aging-related disease onset and death and/or social advantage may confer protection from the effects of aging.

Social structures in human populations are complex and multifaceted, including structural inequities and discrimination, and these factors can vary across cultures. Thus, it can be difficult to measure how social adversity “gets under the skin” in humans to affect immune and overall health. Rhesus macaques (*Macaca mulatta*), a non-human primate, are an established animal model that exhibits aging trajectories similar to humans, such as decreases in mobility, but compressed into a lifespan 3-4x times shorter [23]. Aging parallels are also manifested at the molecular level: rhesus macaques and humans show similar age-related alterations in immune cell DNA methylation and gene expression [24]. Rhesus macaques also share many social factors with humans including the expression of affiliative behaviors, despotism, among other behaviors [25], making them an ideal animal model for translational aging research.

In rhesus macaques, exposure to social adversity can be captured by measures of social status (i.e., dominance rank). Social status is acquired differently for males and females [26–27]: females inherit status from their mothers, while males typically disperse from their natal group and enter a new group where they acquire status through a combination of “queuing” and physical contests [28–29]. Similar to humans, social status in macaques patterns access to resources, and can impact health and lifespan [30–32]. For instance, low status macaques experience more conspecific aggression and are therefore more likely to be injured [33], and high status female macaques can live longer than those with lower status [34]. In addition, social status affects immunity in rhesus macaques; one experimental study showed that social status predicts gene expression patterns in peripheral blood mononuclear cells [35].

Here, we characterized age-related variation in immune cell types, as well as the influence of social adversity and sex on immune cell composition. To do so, we studied a free-ranging population of rhesus macaques living on the island of Cayo Santiago, Puerto Rico where we were able to simultaneously document age, sex, and social status in a semi-naturalistic social setting with minimal human intervention [36].

## Methods

### Study population

Cayo Santiago is an island located off the southeastern coast of Puerto Rico inhabited by approximately 1,600 rhesus macaques. The population is managed by the Caribbean Primate Research Center (CPRC) and is the oldest primate field station in the world [37]. Cayo Santiago provides unique research opportunities for behavioral, physiological, demographic, morphological and genomic studies. The Cayo Santiago Field Station has a minimal intervention policy, which means that the animals are not managed medically or reproductively. There are no predators on the island, and senescent phenotypes are commonly observed in this population [24,38–41]. The animals are direct descendants of rhesus macaques introduced from India in 1938; since 1957 these animals have been continuously monitored [42]. The animals are identified with tattoos and ear notches, and demographic data such as age, sex and pedigree have been collected for decades. During the annual capture and release period, researchers have the opportunity to collect biological and morphological samples with the assistance of CPRC veterinary staff. For the past 15 years, the Cayo Biobank Research Unit has collected detailed behavioral data to combine with the biological samples collected each year. In combination, these data provide the opportunity to test the relationships between the social environment, immune function, and aging.

### Blood sampling

We collected whole blood from sedated rhesus macaques over three capture and release periods (n=96 in October - December 2019, n=153 in October 2020 - February 2021 and n=120 in October 2021 - February 2022). Samples were collected in 6ml K2 EDTA tubes (Beckton, Dickson and Company, cat #367899). We collected a total of 369 samples (200 from males, 169 from females) from 230 unique individuals (113 males, 117 females; i.e., some animals were sampled across multiple years of the study), spanning the natural lifespan of macaques on Cayo Santiago (mean age = 11.8 years, range 0-28 years; **Figure 1A and 1B**). Fresh blood samples were transported at 4°C to the University of Puerto Rico Medical Sciences campus where flow cytometric analysis was performed within 6 hours of sample collection.

**Figure 1:**
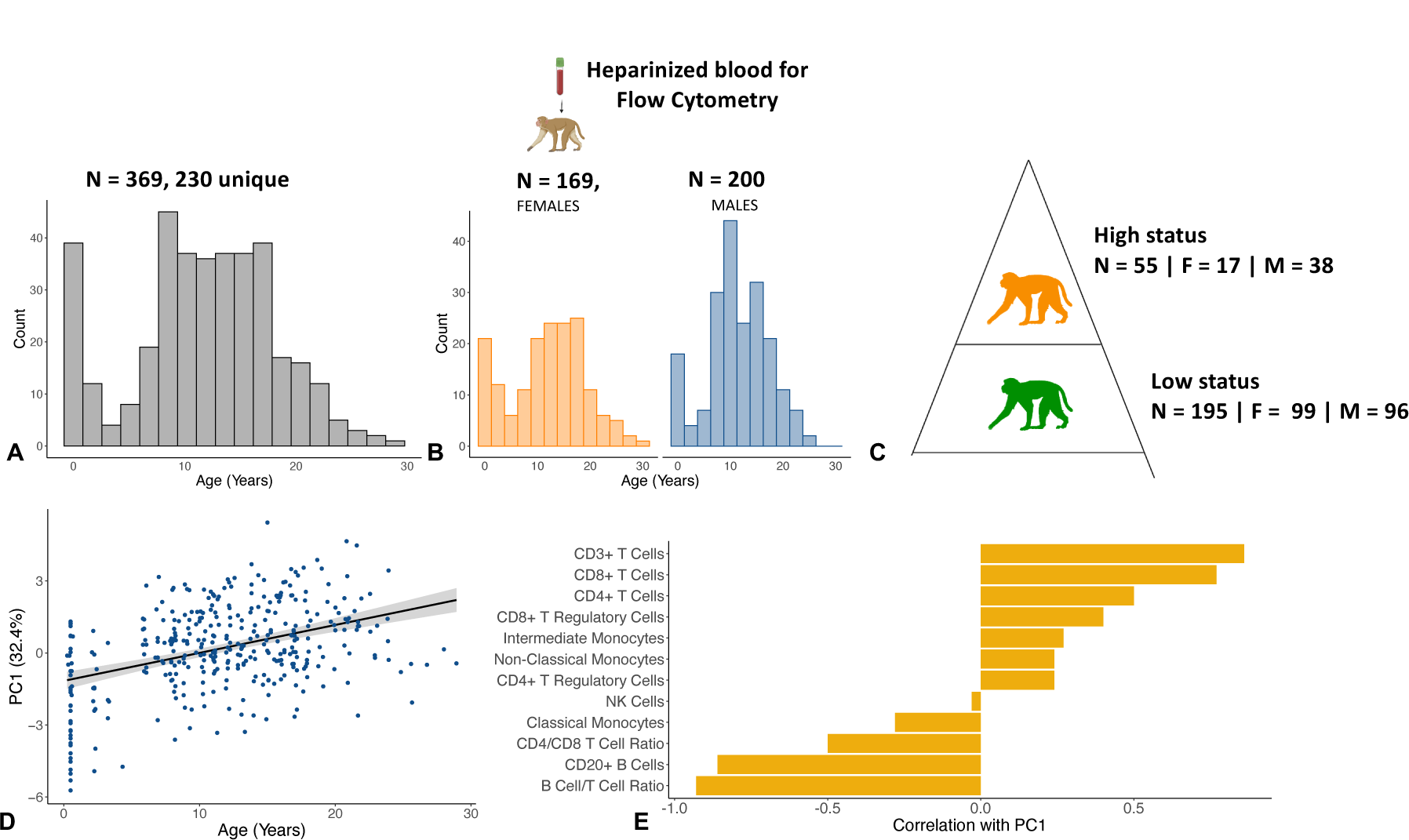
Sample collection and demographics. **A)** We collected 369 whole blood samples from 230 unique individuals across three years, and quantified immune cell proportions using flow cytometry. **B)** The dataset was roughly balanced between males and females and captured the entire natural lifespan of macaques in this population. **C)** We calculated social status by assigning dominance ranks to 250 samples using observational data collected in the year before each sample was collected. Animals were assigned to one of three dominance ranks: high, medium, and low. The social status dataset is a subset of the original age dataset because behavioral data were not available for all study animals (i.e., it is not collected for infants and juveniles). **D)** PC1 of immune cell compositions is significantly associated with age (β_PC1_ = 0.14, FDR = 7.2 x 10^-18^). **E)** The T cell compartment is positively associated with PC1 (and thus age), while the B cell compartment is negatively associated with PC1 of immune cell composition.

### Antibodies and flow cytometric analysis

An 8-panel antibody cocktail previously validated in rhesus macaques [43–46], consisting of the following antibodies, was used: CD20-PacBlue/Clone 2H7 (Biolegend), CD3-PerCP/Clone SP34-2 (BD), CD4-APC/Clone L200 (BD), CD8-Viogreen/Clone BW135/80 (Miltenyi), CD25-PE/Clone 4E3 (Miltenyi), CD14-FITC/Clone M5E2 (BD), CD16-PEVio770/Clone REA423 (Miltenyi), HLA-DR-APCVio770/Clone REA805 (Miltenyi).

We performed phenotypic characterization of rhesus macaque peripheral blood mononuclear cells (PBMCs) using multicolor flow cytometry with direct immunofluorescence (View **S. Figure 1** for gating strategy and **Table S1** for Ab panel) on all 369 animals. Aliquots of 150 μl of heparinized whole blood were incubated with a mix of the antibodies described for 30 minutes at 25°C (room temperature). After incubation, the red blood cells were fixed and lysed with BD FACS fix and lyse solution (Cat #349202). Cells were washed twice using PBS containing 0.05% BSA at 1,700 RPM for 5 minutes and processed in a MACSQuant Analyzer 10 flow cytometer (Miltenyi Biotec, CA).

Lymphocytes and monocytes were gated based on their characteristic forward scatter (measures cells based on their size) and side scatter (measures cells based on their granularity) patterns. Lymphocytes were then further subdivided according to their cell surface markers. Natural killer (NK) cells were defined as the CD3- and CD16+ population; B cells were defined as CD20+ population and T cells as the CD3+ population. We further subdivided T cells from the CD3+ gate into CD4+ and CD8+ populations. CD4+CD25+ and CD8+CD25+ T regulatory cells were further gated from the CD4+ and CD8+ gates. Monocytes were gated based on the combined expression of the HLA-DR/CD14 markers for classical monocytes, HLA-DR/CD16 markers for non-classical monocytes, and HLA-DR/CD14/CD16 for intermediate monocytes (**S. Figure 2**). Flow cytometry gating was performed using Flowjo version 10.7.1 (FlowJo LLC Ashland, OR).

**Figure 2:**
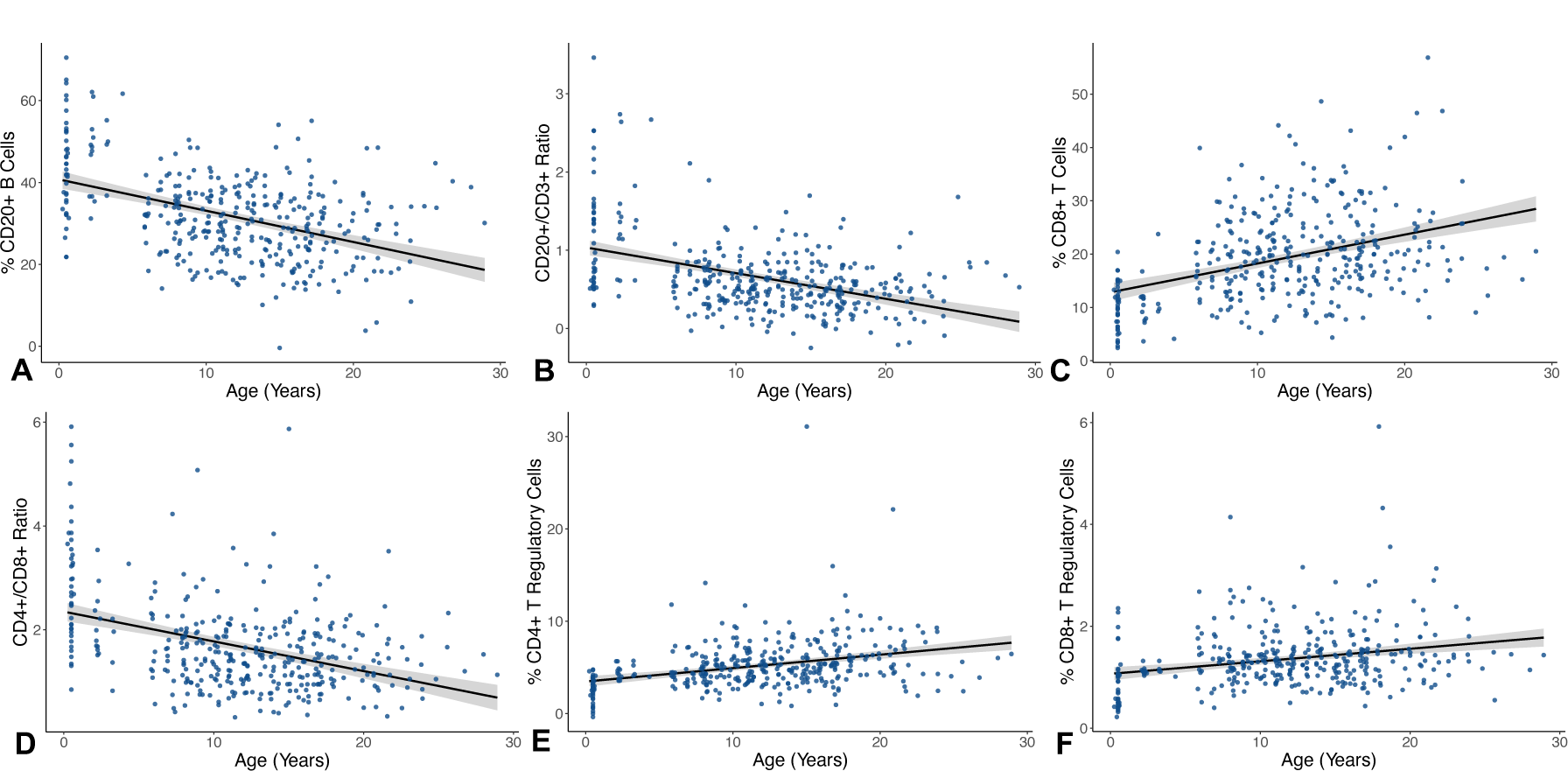
Age-associated differences in adaptive immune cell proportions. **A)** CD20+ B cells (β_CD20_ = −0.83 ± 0.09, FDR = 1.3 x 10^-16^) proportions and **B)** CD20+/CD3+ ratio (β_CD20:CD3_ = −0.04 ± 0.004, FDR = 4.9 x 10^-15^) are lower in older individuals. **C)** CD8+ Cytotoxic T cells (β_CD8_ = 0.60 ± 0.07, FDR = 3.2 x 10^-14^) are higher in older individuals, while the **D)** CD4+/CD8+ T cell ratio (β_CD4:CD8_ = −0.06 ± 0.008, FDR = 4.3 x 10^-14^) is lower in older individuals. **E)** CD4+ T regulatory cells (β = 0.16 ± 0.02, FDR = 3.6 x 10^-10^) and **F)** CD8+ T regulatory cells are higher in older individuals compared to younger individuals (β = 0.03 ± 0.005, FDR = 1.8 x 10^-6^), possibly because of higher baseline levels of inflammation (i.e., inflammaging).

To obtain an accurate representation of the proportions of cell types, we counted only the stained events of the cells of interest and calculated their proportions based on the subsets of lymphocytes and monocytes. To calculate cell ratios, such as CD20 + B cell to CD3+ T cell ratio (CD20+/CD3+ ratio) and CD4+ T cell to CD8+ T cell ratio (CD4+/CD8+ ratio), we divided the calculated proportion of these cell types in each individual sample (e.g., CD20+ B cell/ CD3+ T cell and CD4+/CD8+ respectively). (Calculations detailed in **Table S2**).

### Quantification of social adversity (social status)

We quantified the social status of a subset of animals for which we had behavioral data (**Figure 1C,** n = 250 total samples, 134 from males and 116 from females & n = 145 unique individuals, 73 males and 72 females). We calculated social status (i.e., dominance rank) using the outcomes of win-loss dominance interactions between pairs of same sex adult groupmates. Because of the different routes through which males and females acquire their status, we quantified social status separately for males and females within each group for each year of the study [47–48]. Observations of adult animals (older than 6 years) were collected from two different social groups, groups V and F, in the year prior to sample collection. In 2019 and 2021, behavioral data were collected using a previously established 10-minute focal sampling protocol [26]. Briefly, the protocol consisted of recording state behaviors (e.g., resting, feeding) and agonistic encounters, which included recording the identity of the focal animal and their social partner. Win-loss agonistic interactions included threat and submissive behaviors, along with contact and non-contact aggression. In 2019 and 2021 we also collected additional agonistic interactions *ad-libitum*. In 2020, all agonistic interactions were collected *ad-libitum* due to restrictions imposed on behavioral data collection due to the COVID-19 pandemic. In all years, we used known maternal relatedness to settle behavioral gaps in the female hierarchy [49]. To control for variation in group size, social status (i.e., dominance rank) was quantified as the percentage of same-sex adults that an animal outranked in their group. For all analyses, we grouped animals into one of two social status categories: high-rank (>80% of same-sex adults outranked) and low-rank (< 79% of same-sex adults outranked)[50].

### Statistical analysis

All statistical analyses were performed using R statistical software R version 4.2.3 [51]. First, we performed a principal component analysis of the cell composition data for all samples (n = 369, 230 unique individuals) using the *prcomp* function of the *stats* package. Next, we employed a linear additive mixed-effects approach, using the *lmer* function in the *lmerTest* package to run sample projections on principal components as a function of age (in years), sex, and sample period - to control for the technical variation in the flow cytometer lasers, which changed over the sampling years (*model 1* - **Table S3**) and individual ID as a random effect. We also modeled sample projections on principal components as a function of the interaction between age and sex (age*sex) and sampling period - which will ultimately allow us to identify possible sex-dependent associations with age - and individual ID as a random effect (*model 2* - **Table S3**).

To evaluate each cell type at a more granular level, we employed the same additive linear mixed-effects to test the proportion of each cell type and certain cell type ratios (e.g., CD4+/CD8+) as a function of age, sex, and sample period with individual ID as a random effect (*model 3* - **Table S3**). Finally, we tested the proportion of each cell type and certain cell type ratios (e.g., CD4+/CD8+) as a function of the interaction between age and sex (age*sex) and sampling period with individual ID as a random effect (*model 4* - **Table S3**).

For the subset of samples in which information on social status was available (n = 250 total samples, 145 unique individuals), we tested for the additive effect of principal component projections as a function of social status, age, sex and sample period, with individual ID and social group as a random effect (*model 5* - **Table S3**). We also tested for the principal component interaction between status and age (status*age) and between status and sex (status*sex), with individual ID and social group as a random effect (*model 6* - **Table S3**). We then additively tested the proportion of each cell type and certain cell type ratios as a function of social status, age, sex, and sample period (*model 7* - **Table S3**). To test whether the relationship between the proportion of cell types and social status depended on sociodemographic variables, we tested the interaction between: social status and age (status*age) and for social status and sex (status*sex, *model 8* - **Table S3**). Furthermore, since we identified interactions between social status and sex, and since social status is acquired differently for male and females in rhesus macaques, we fitted a separate model for males and females to test if there was a main effect of social status within each sex on the proportions of different cell types. Age and sample period were also included in the model to control for these covariates (*model 9* - **Table S3**).

The linear models and sample sizes for each are summarized in **Table S3.** For every predictor variable in the full (n = 369) and status (n = 250) datasets, we corrected for multiple hypothesis tests using the Benjamini Hochberg FDR approach and considered significant associates at FDR < 0.10. Because Model 7 was only performed in cell types that showed a significant interaction between sex and social status (and not all the cell types measured), we did not correct for multiple testing in this model.

## Results

### Macaques exhibit age-related variation in immune cell composition and inflammation

Age was significantly positively associated with the first principal component (PC) of immune cell composition (*model 1* - β_PC1 age_ = 0.14, FDR = 7.2 x 10^-18^, **Figure 1D)**. This first PC, which explained 32.4% of the variance in cell composition across all samples, was associated with higher proportions of inflammation-associated cell types, such as cytotoxic T cells and regulatory T cells, and lower proportions of cells involved in pathogen clearance, including CD20+ B cells and classical monocytes (**Figure 1E, Table S4).** Thus, older animals exhibited a pattern of greater inflammation and immunosenescence than younger individuals did.

We then conducted a more granular analysis of the factors associated with the proportions of individual cell types. Age was significantly associated with signatures of immunosenescence, including a decline in adaptive immune cells. This was largely driven by lower proportions of CD20+ B cells in older individuals (*model 3* - β_CD20 age_ = −0.83 ± 0.09, FDR = 1.3 x 10^-16^, **Figure 2A**), which resulted in significantly lower CD20+/CD3+ ratios in older individuals (*model 3* - β_CD20:CD3 age_ = −0.04 ± 0.004, FDR = 4.9 x 10^-15^, **Figure 2B**). Age was also associated with higher proportions of inflammation-related cells. The proportion of cytotoxic CD8+ Cytotoxic T cells was significantly higher in older animals (*model 3* - β_CD8 age_ = 0.60 ± 0.07, FDR = 3.2 x 10^-14^, **Figure 2C**), resulting in a strong and significant effect of lower CD4+/CD8+ ratios (*model 3* - β_CD4:CD8 age_ = −0.06 ± 0.008, FDR = 4.3 x 10^-14^, **Figure 2D**) and higher proportions of CD3+ T cells in older individuals (*model 3* - β_CD3_= 0.67 ± 0.11, FDR = 2.2 x 10^-8^, **S. Figure 3)**.

**Figure 3:**
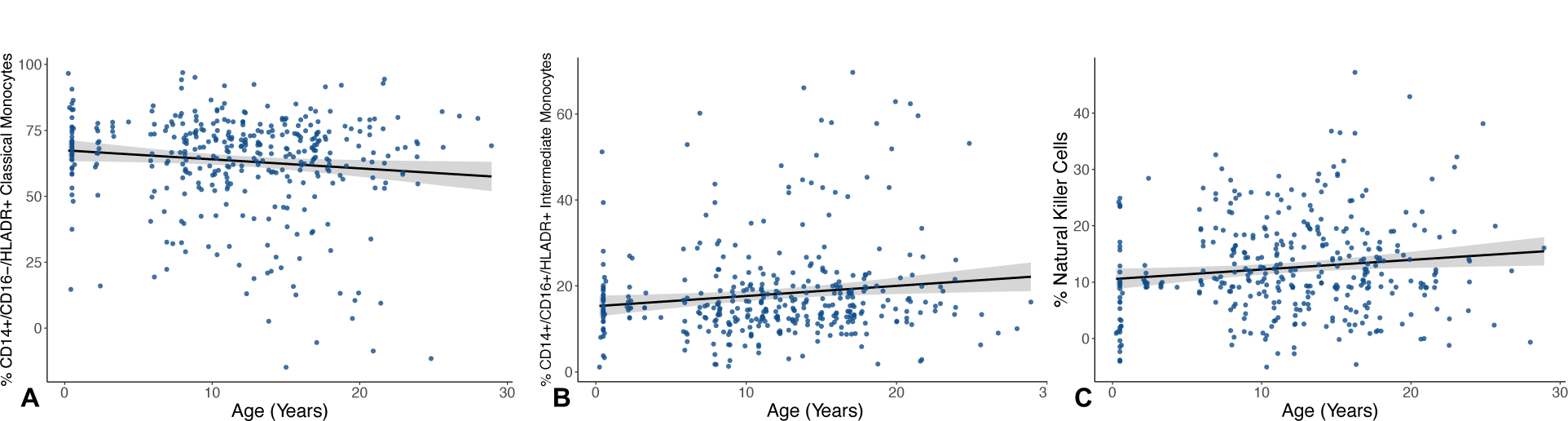
Age is associated with variation in innate immune cell proportions. **A)** CD14+/CD16-/HLADR+ Classical monocytes (β_CD14++_ = −0.31 ± 0.16, FDR = 0.07) are lower and **B)** CD14+/CD16+/HLADR+ intermediate monocytes (β_CD14+CD16+_ = 0.21 ± 0.09, FDR = 0.04) are higher in older individuals, while **C)** CD16+ NK cells (β_NK_ = 0.17 ± 0.07, FDR = 0.03) are higher in older individuals.

Next, we examined the less abundant but immunologically important regulatory CD8+ and CD4+ T cell populations (CD25+), which are involved in immune suppression and maintenance of self-tolerance [52] (i.e., the ability to recognize self-antigens). Both CD4+ and CD8+ T regulatory cells were significantly more abundant in older animals (*model 3* - CD3+CD4+CD25+: β _age_= 0.16 ± 0.02, FDR = 3.6 x 10^-10^, **Figure 2E**; *model 3* - CD3+CD8+CD25+: β _age_ = 0.03 ± 0.005, FDR = 1.8 x 10^-6^, **Figure 2F**), suggesting a reduced age-related ability to regulate endogenous and exogenous antigens.

Innate immune cells also showed significant associations with age. Classical monocytes (HLA-DR+/CD14+/CD16-), which are involved in phagocytosis and extracellular pathogen clearance [53], were lower in older individuals (*model 3* - β_CD14++ age_ = −0.31 ± 0.16, FDR = 0.07, **Figure 3A**), while intermediate monocytes (HLA-DR+/CD14+/CD16+), involved in immune cell recruitment and proinflammatory cytokine secretion [53], were higher in older individuals (*model 3* - β_CD14+CD16+ age_ = 0.21 ± 0.09, FDR = 0.04, **Figure 3B)**. The proportion of CD16+CD3-NK cells – which have a similar role to CD8+ Cytotoxic T cells presenting natural cytotoxicity but are not antigen specific – was also significantly higher in older individuals (*model 3* - β_NK age_ = 0.17 ± 0.07, FDR = 0.03, **Figure 3C**). Together, these results indicate that older individuals show a decrease in adaptive immunity along with an increase in inflammation-related innate immune cells compared to younger individuals, potentially disrupting a “healthy” homeostatic immune system.

We did not observe statistically significant main effects of sex (*model 3* in *Methods*) or a sex-age interaction (*model 4* in *Methods*) on immune cell proportions. Nevertheless, a trend toward sex differences was observed in both the proportions of CD8+ Cytotoxic T cells (*model 3* - β_CD8 sex_ = 2.19 ± 0.95, FDR = 0.14) and in the CD4+/CD8+ ratio (*model 3* - β_CD4:CD8 sex_ = −0.24 ± 0.09, FDR = 0.14, **S. Figure 4**), with males having a higher proportion of CD8+ Cytotoxic T cells compared to females, and females having a higher CD4+/CD8+ ratio compared to males, suggesting a stronger adaptive immune response in females, which, in part, is generated by CD4+ T helper cells.

**Figure 4:**
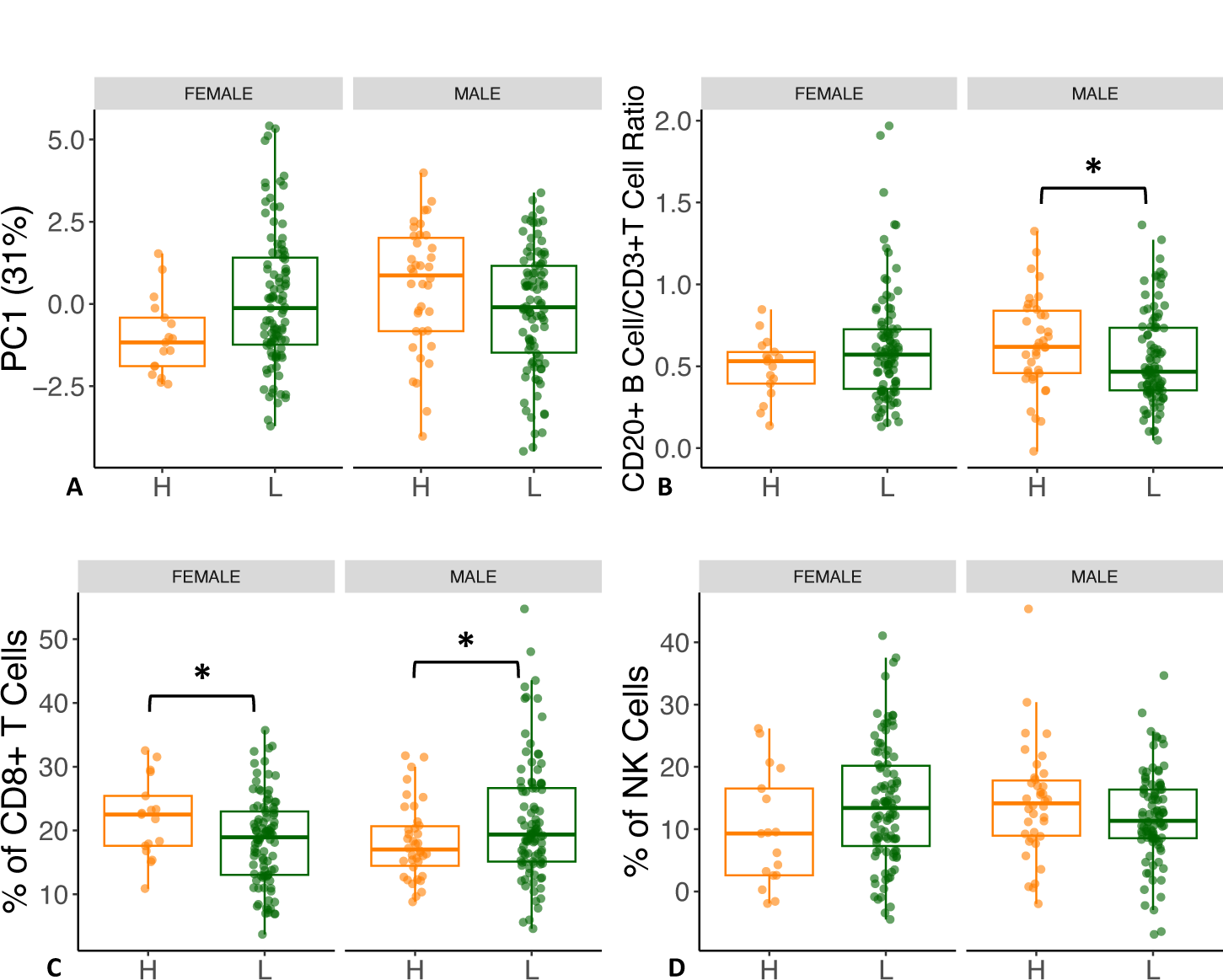
Sex and social status interact to impact immune cell composition. **A)** PC1 (31% of the variation in the dataset, β_PC1-sex*status_ = −1.7, FDR = 0.03) recapitulates the interaction between sex and social status. **B)** Interaction in the CD20+/CD3+ ratio (β_CD20/CD3 ratio-sex*status_ = −0.25 ± 0.05, FDR = 0.06) shows that this ratio is higher with higher social status in males while it is lower with higher social status in females; within-sex analysis show that high social status males have a significantly higher ratio than low social status males (β_CD20/CD3 ratio-males/status_ = −1.3 ± 0.06, p = 0.04). **C)** CD8+ Cytotoxic T cells show an interaction (β_CD8-sex*status_ = 8.3 ± 2.94, FDR = 0.04) where this cell type is higher in lower social status in males, while it is higher in high social status females. Within-sex analysis of CD8+ T cells showed low social status males had significantly higher proportions of this cell type than did high social status males (β_CD8-males/status_ = 4 ± 1.8, p = 0.03), while the opposite effect was observed in females (β_CD8-females/status_ = −4.8 ± 2.1, p = 0.03). **D)** Interaction in the proportion of NK Cells (β_NK-sex*status_ = −7.9 ± 3.2, FDR = 0.05), with high social status males showing higher proportions, while high social status females show lower proportions.

### Social status and immune cell composition

There was a significant interaction between social status and sex on PC1 (31% of the variance in cell composition across all samples) of immune cell composition (*model 6* - β_PC1-sex*status_ = −1.7 ± 0.63, FDR = 0.04, **Figure 4A)**, documenting the sex-dependent impact of social status on immunity.

When modeling males and females together in an additive modeling framework, we found no significant effects of social status on immune cell proportions (all FDR > 0.10, *model 7* in *Methods*), or between the interaction between social status and age (*model 8* in *Methods*). However, we found many significant interactions between social status and sex on immune cell composition (*model 8* in *Methods*). Because social status is acquired differently for male and female rhesus macaques, we also carried out post-hoc analyses of the social status effects within each sex separately (*model 9* in *Methods*).

The CD20+/CD3+ ratio interacted between social status and sex such that it was higher in males of high social status, but lower in females with high social status (*model 8-* β_CD20/CD3 ratio-sex*status_ = −0.26 ± 0.11, FDR = 0.06, **Figure 4B)**, both part of the adaptive immune system. This difference in the CD20+/CD3+ ratio seems to be partially driven by interaction between sex and social status on the proportion of CD3+ T cells (*model 8-* β_CD3-sex*status_ = 11.8 ± 4.42, FDR = 0.04, **S. Figure 5)**, such that CD3+ T cells were higher in low social status males, but this pattern was flipped in females. There was a within-sex main effect of social status in males in the CD20+/CD3+ ratio, with high social status males having a significantly higher ratio than low social status males did (*model 9-* β_CD20/CD3 ratio-males/status_ = −1.3 ± 0.06, p = 0.04, **Figure 4B**). No significant effect of social status on the CD20+/CD3+ ratio was found in females (*model 9-* β_CD20/CD3 ratio-females/status_ = 0.10 ± 0.13, p = 0.19).

Additionally, there was a significant interaction between social status and sex in the proportion of CD8+ Cytotoxic T cells (*model 8-* β_CD8-sex*status_ = 8.3 ± 2.94, FDR = 0.04, **Figure 4C)**, which were higher in high social status males, but this relationship again flipped females. Our within-sex analysis revealed a significant main effect of CD8+ Cytotoxic T cells, in which low social status males had significantly higher proportions of this cell type than did high social status males (*model 9-* β_CD8-males/status_ = 4 ± 1.8, p = 0.03, **Figure 4C**). The opposite main effect was observed in females, in which high status females had significantly higher proportions of CD8+ Cytotoxic T cells than did low social status females (*model 9-* β_CD8-females/status_ = −4.8 ± 2.1, p = 0.03, **Figure 4C**).

In the innate arm of the immune system, we detected an interaction between social status and sex on the proportion of CD3-CD16+ NK cells (*model 8-* β_NK-sex*status_ = −7.9 ± 3.2, FDR = 0.05, **Figure 4D)**, where the proportion was lower in males of low status, but higher in females of low status. Social status approached significance in the proportions of CD3-CD16+ NK cells in females (*model 9-* β_NK-females/status_ = 5.6 ± 3, p = 0.06, **Figure 4D**), in which low social status females displayed higher proportions of this cell type compared to high social status females. We found no significant main effect of social status on CD3-CD16+ NK cells in males (*model 9-* β_NK-males/status_ = −2.27 ± 3.4, p = 0.18).

## Discussion

We examined how social status, age, and sex were related to immune cell distributions in a large sample of adult rhesus macaques living in semi-natural conditions. Overall, we found strong and consistent signatures of age-related immune cell dysregulation. We also identified significant links between social status and sex in cells of innate and adaptive arms of the immune system. Together, this variation is likely to influence immune responses to pathogenic challenges as well as the development of inflammation-related diseases.

Overall, macaques exhibited age-related differences in immune cells similar to those observed in humans, including declines in lymphocytes [54]. Here, we also identified more specific cell types with age-related differences. We detected lower proportions of CD20+ B cells at older ages, which may reflect immunosenescence, as these cells are responsible for antibody production, pathogen clearance, and are key cells in the generation of immune memory. Further, a key factor underlying the limited efficacy of vaccines in older individuals is a weakened B cell response [55]; B cells have also been associated with protection against certain types of cancer, such as lung cancer [56].

Similar to two other studies in captive macaques, we found higher CD8+ T cell proportions at older ages [67–58]. Notably, this differs from findings in humans, where both CD8+ Cytotoxic T cells and their effector responses (i.e., stimulus responsiveness) exhibit lower proportions at older ages [59]. It is possible that this discrepancy is only reflected in the overall CD8+ cytotoxic T cell pool, as it has been reported that certain CD8+ T cell subsets – such as memory subsets – increase in proportion and efficacy with age [60]. Alternatively, given that CD8+ T cell subsets have been associated with inflammation and ‘inflammaging’ [61], there is a possibility that higher overall CD8+ Cytotoxic T cell pool in rhesus macaques is indicative of higher levels of inflammation. The age-related reduction in CD4+/CD8+ ratio corroborates this hypothesis. In support of increased inflammation with age, we found that older animals had significantly higher CD3-CD16+ NK cell proportions in our dataset. Similar to CD8+ Cytotoxic T cells, CD3-CD16+ NK cells respond to intracellular pathogens, secrete multiple proinflammatory mediators, and are crucial during tumor surveillance and signaling [62]. The higher proportions of CD3-CD16+ NK cells predict a higher incidence of inflammation and/or tissue injury in the older population, which is commonly observed in the Cayo Santiago macaque population [33]. As expected, CD4+CD25+ T regulatory cells as well as CD8+CD25+ T regulatory cells were associated with age, indicating higher levels of inflammation in older individuals [63]. These results, together with lower levels of CD20+ B cell proportions and higher levels of CD8+ T cell and CD3-CD16+ NK cell proportions, further support the hypothesis that the adaptive immune response in rhesus macaques decreases with age and inflammation-related cell types increase (i.e., ‘inflammaging’). Taken together, these alterations may drive biological and physiological decline that likely increases the risk of morbidity and mortality in macaques, as it does in humans.

Monocyte proportions also varied with age. Specifically, we found fewer CD14+ classical monocytes in older animals. These cells are phagocytic cells that ingest pathogens that they encounter [64]. This age-related reduction may indicate a reduction in phagocytosis (ingestion of pathogens by classical monocytes) and thus can possibly increase infections in older individuals. In addition, older individuals had higher proportions of CD14+/CD16+ intermediate monocytes, which are strongly associated with inflammation [65]. For instance, an increase in this cell type has been linked to disorders such as chronic kidney disease [66]. The decrease in classical monocytes, together with an increase in intermediate monocytes, represents yet another signature of immunosenescence and inflammaging.

One of the strengths of our study system was the ability to quantify social adversity, operationalized as social status, and test if and how social status influenced immune variation and whether the effects of status varied with age and/or sex. We found no main effect of social status on the proportion of immune cell types. Also, there was no interaction between social status and age on the proportions of immune cell types. This result was contrary to our expectations because we expected low social status individuals to experience more variation in immune cell types with increasing age. Nevertheless, we found several interactions between social status and sex, as well as a within-sex main effect of social status on immune cell composition, possibly reflecting the different pathways through which social status is acquired in males and females and thus highlighting the fact that different sexes experience social adversity differently across the hierarchy.

The interaction of social status and sex influenced adaptive and innate immune cell types such as CD20+/CD3+ ratio and CD8+ T cells and CD3-CD16+ NK cells, where the proportions of these cell types associated with status depended on the sex of the individual. In humans, social stressors, such as lower socioeconomic status and lower subjective social status, can affect cytokine release and inflammatory responses in peripheral blood mononuclear cells in a sex-dependent manner [67–68]. However, studies in humans that have looked at the interaction between social stressors (such as socioeconomic status) and sex on immune cell proportions have found no significant interaction between these covariates [22], thus making our study unique in reporting sex-dependent effects of social status.

We also found a significant main effect of social status in the within-sex analysis on the CD20+ B cell/CD3+ ratio, with high social status males having significantly higher ratios than low status males. The decrease in the CD20+/CD3+ ratio seems to be driven by a decrease in the proportions of CD3+ T cells in low social status males (**S. Figure 5**) compared to high status males. Decreases in this cell type have been associated with decreases in cell-mediated immunity to bacteria and viruses [69–70], potentially showing that the T cell response in macaques is negatively affected by low social status. In addition, CD8+ T cell proportions were higher in low social status males compared to high social status macaques. Few prior studies have assessed sociality-related immune cell differences in male rhesus [71], likely because of ethical and husbandry constraints, such as aggressive behavior between males. One study in male Barbary macaques (*Macaca sylvanus*) reported that males with strong social bonds had lower levels of fecal glucocorticoids [72], which is typically associated with reduced inflammation [73]. Additionally, studies in cynomolgus macaques (*Macaca fascicularis*) have shown that low social status males had a higher probability of being infected with a virus than did high social status macaques [74]. These findings should be taken with caution, however, as other studies of macaques (rhesus and other species) found no differences in infection rate or immune responses between high and low status males [75–76]. Although there are currently no data associating social status (or other social stressors) with CD8+ T cells in rhesus macaques, there are reports in other species that CD8+ T cells can mediate the release of proinflammatory cytokines during stressful conditions [77]. Our finding of higher proportions of CD8+ T cells in low social status macaques might indicate higher levels of baseline cytotoxic T cell activation, potentially affecting the CD4+ T cell response. Testing this idea will require methods such as cytokine analysis or next generation sequencing.

There was also a main effect of social status on the proportion of CD8+ T cells in females, but in contrast to males, high social status females had significantly higher proportions of this cell type compared to low social status females. One study also reported lower proportions of CD8+ T cells in low social status in non-free-ranging female rhesus macaques [34]. Given that females tend to have lower proportions of CD8+ T cells than males regardless of age [78–80], a lower proportion of this cell type in low social status females might indicate lower cytotoxic immunity at baseline. Female social status also had a main effect on the proportion of CD3-CD16+ NK cells (associated with immune surveillance, inflammation and innate responses), with low social status females having significantly higher proportions of this cell type than high social status females. Although a prior study found that the proportion of CD3-CD16+ NK cells did not vary with social status in female rhesus macaques, it did find that this cell type was the most sensitive to social status. Specifically, low social status females showed patterns of gene expression consistent with a proinflammatory phenotype in this cell type in response to lipopolysaccharide [81]. These results highlight that low status female rhesus macaque may experience higher levels of basal inflammation, consistent with other studies in this species [81–83].

In conclusion, our results demonstrate that, at the level of circulating immune cell proportions, macaques and humans show similar age-related variation in immune cell types. Although we did not detect any significant main effects of sex or sex-age interaction, it is possible that more specific, but unmeasured adaptive immune cells, such as the effector and memory subsets of B cells and T cells, could differ between males and females. In future studies, it will be important to measure other innate immune cell types, such as dendritic cells and granulocytes, since these cell types are critical for antigen presentation and the development of adaptive immune response. We found that the effects of social status differed between males and females, which is likely due to sex-differences in how rhesus macaques obtain social status. Specifically, females inherit their social status, which remains relatively stable throughout their lives, while males queue and occasionally fight to establish and maintain their social status, which may lead to stronger effects of status on immune cell distribution and function. Overall, our study provides detailed insights into the impacts of social and demographic variation on immune cell status in a non-human primate model with unparalleled translatability to humans. Future research should quantify the proportions of these cell types as a function of age using a longitudinal approach, which will require sampling individuals over the course of years. Immune stimulation tests would also be informative by testing whether both the age-associated and status-associated differences in immune cell types translate to immune function.

## Author contributions

M.R.S.R., J.P.H., L.J.N.B., C.A.S., M.J.M, M.L.P., and N.S.-M designed research; M.R.S.R., N.M.R., M.M.W., A.D.N.-D, P.P., M.A.P.-F., E.R.S., E.B.C., J.E.N.-D., D.P., A.R.L., M.J.M. and CBRU performed research; M.R.S.R and N.S.-M. analyzed data; and M.S.R. and N.S.-M. wrote the manuscript. All authors reviewed and revised the manuscript. CBRU members: Lauren J.N. Brent, Jampes P. Higham, Noah Snyder-Mackler, Michael J. Montague, Michael L. Platt, Melween I. Martinez, Susan C. Antón, Amanda D. Melin, Jérôme Sallet The authors declare no competing interest.

## Acknowledgements

Part of Figure 1, panel A, was created with BioRender.com. We thank the Caribbean Primate Research Center for their help in collecting data, especially Giselle Carabllo Cruz, and Nahiri Rivera Barreto; Crisanta Serrano and Stephanie Dorta for laboratory advice; Corbin Johnson, Laura Newman, Alice Baniel, India Schneider-Crease, Kenneth Chiou, Trisha Zintel, and other members of the Snyder-Mackler, Sariol, Higham, Platt, and Brent labs for helpful discussions during the course of this work.

## Ethical note

This work was approved by the Institutional Animal Care and Use Committees of the University of Puerto Rico, Medical Sciences Campus (IACUC Number: A400117).

The authors declare no competing interest.

