## Supplementary Material for "Age, sex, and social environmental effects on immune cell composition in a free-ranging non-human primate"

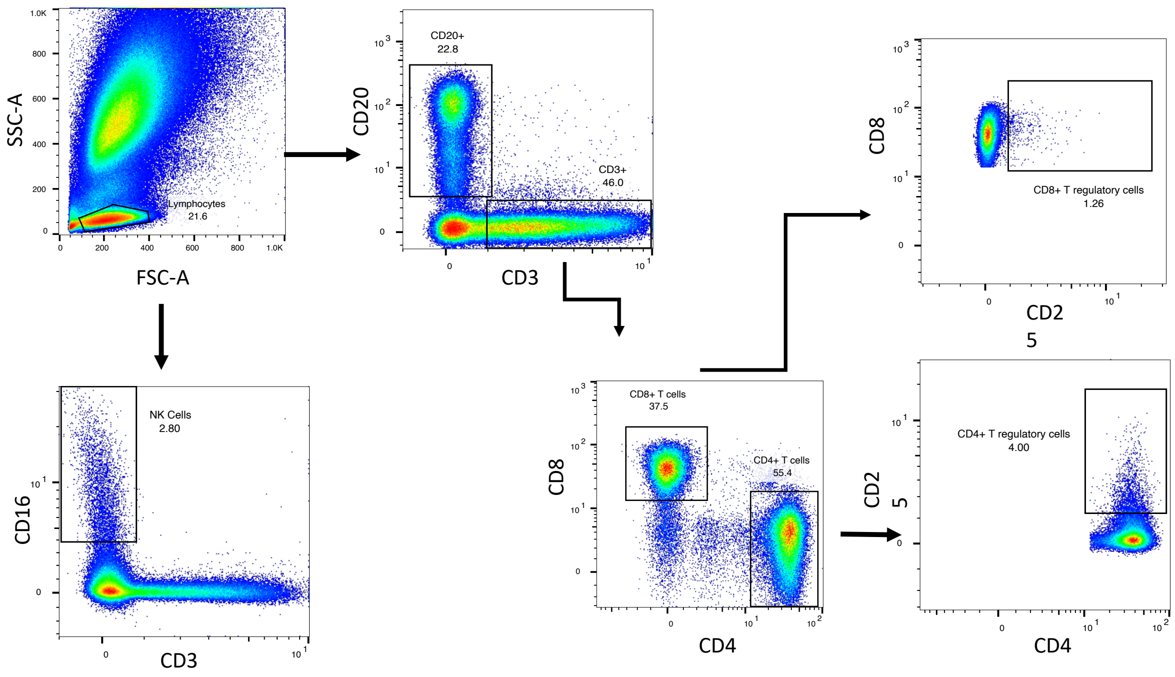


**Supplementary Figure 1. Lymphocyte gating strategy.** Lymphocytes were gated based on their characteristic forward and side scatter patterns (FSC, SSC). B cells were identified as CD20+/CD3--; NK cells were gated as the CD16+/CD3- population; CD4 + and CD8+ T cells were defined as CD3+ CD4+ and CD3+ CD8+, respectively.

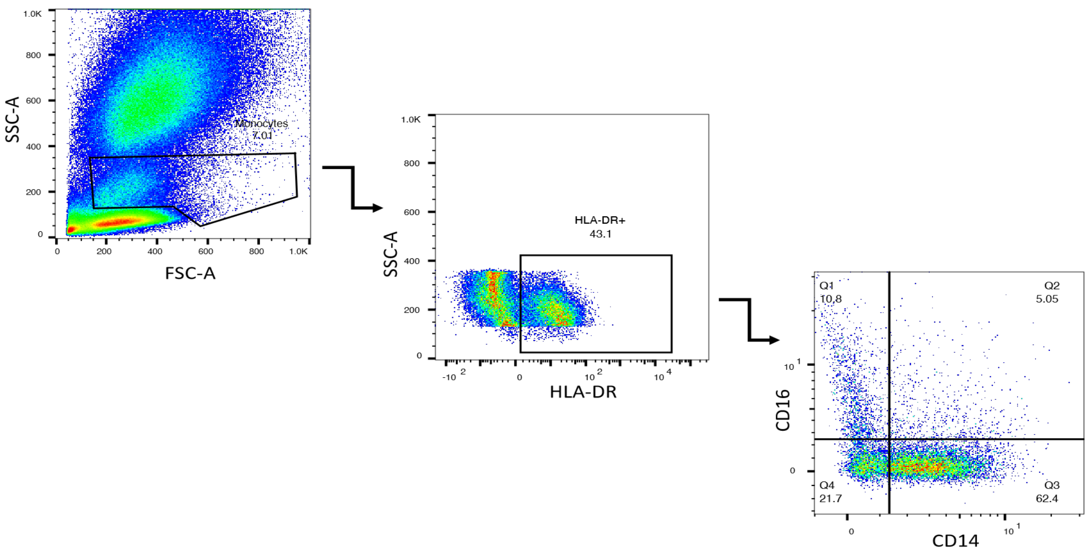


**Supplementary Figure 2. Monocyte gating strategy.** Monocytes were gated based on their characteristic forward and side scatter patterns (FSC, SSC). The monocyte population was then selected gating on the HLA-DR+ population. Classical monocytes were gated based on the expression of CD14+, non-Classical monocytes were gated based on the expression of CD16+, and intermediate monocytes were gated based on the combined expression of both CD14+ and CD16+.

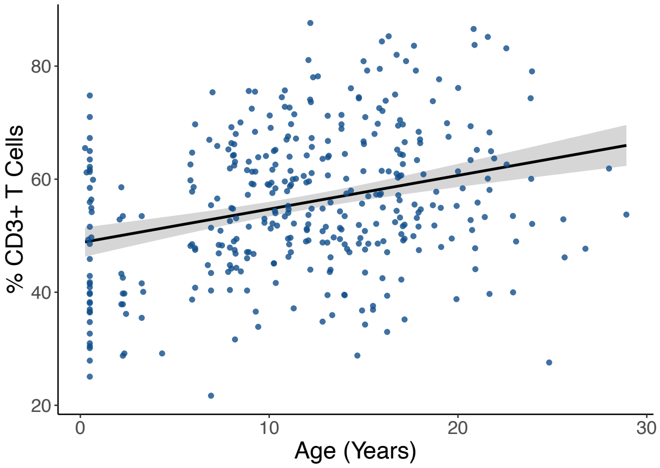


**Supplementary Figure 3.** CD3+ T cell proportions are greater ($\beta_{CD3}$= 0.67 ± 0.11, FDR = $2.2x10^{-8}$) with age.

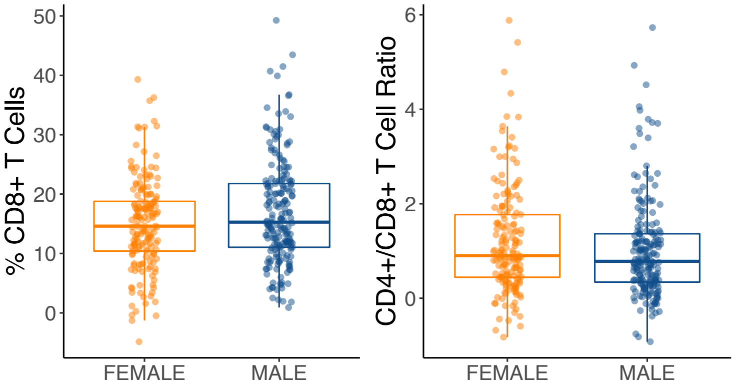


**Supplementary Figure 4.** **A)** Males have, on average, significantly higher proportions of CD8+ T cells ($\beta_{CD8\ sex}$ = 2.19 ± 0.95, FDR = 0.14) and **B)** females have a higher CD4+/CD8+ T cells ratio ($\beta_{CD4:CD8\ sex}$ = -0.24 ± 0.09, FDR = 0.14).

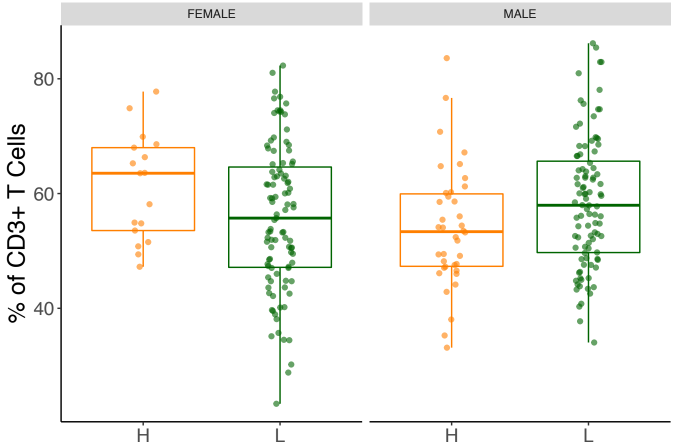


**Supplementary Figure 5.** Interactions between sex and social status where CD3+ T cells proportions are higher ($\beta_{CD3\text{-}sex*status}$ = 11.8 ± 4.42, FDR = 0.04) in high social status females but lower in high social status males.

| Antibody | Dye | Clone | Co. | Cat.# |
|---|---|---|---|---|
| CD3 | PerCP | SP34-2 | BD | 552851 |
| CD20 | Pac Blue | 2H7 | BioLegend | 302328 |
| CD14 | FITC | M5E2 | BD | 555397 |
| CD16 | PE-Vio770 | REA 423 | Miltenyi | 130-113-394 |
| CD4 | APC | L200 | BD | 551980 |
| CD8 | VioGreen | BW135/80 | Miltenyi | 130-113-164 |
| CD25 | PE | 4E 3 | Miltenyi | 130-113-282 |
| HLA-DR | APC-Vio770 | REA 805 | Miltenyi | 130-111-792 |

**Supplementary Table 1.** Antibody panel for Immunophenotyping. All antibodies information is available in the antibody registry page (https://antibodyregistry.org/) and they have been validated in rhesus macaques.

| Cell type | Formula |
| --- | --- |
| B cells | $\frac{(CD20+\ B\ Cells)}{(CD20+B\ Cells)+(CD3+T\ Cells)+(CD16+CD3-NK\ Cells)}$ X 100 |
| T cells | $\frac{(CD3+T\ Cells)}{(CD20+B\ Cells)+(CD3+T\ Cells)+(CD16+CD3-NK\ Cells)}$ X 100 |
| Natural Killer cells | $\frac{(CD16+CD3-\ NK\ Cells)}{(CD20+B\ Cells)+(CD3+T\ Cells)+(CD16+CD3-NK\ Cells)}$ X 100 |
| CD4+ T cells | $\frac{(CD3+CD4+\ T\ Cells)}{(CD20+B\ Cells)+(CD3+T\ Cells)+(CD16+CD3-NK\ Cells)}$ X 100 |
| CD8+ T cells | $\frac{(CD3+CD8+\ T\ Cells)}{(CD20+B\ Cells)+(CD3+T\ Cells)+(CD16+CD3-NK\ Cells)}$ X 100 |
| CD4+ T regulatory cells | $\frac{(CD3+CD4+CD25+\ B\ Cells)}{(CD20+B\ Cells)+(CD3+T\ Cells)+(CD16+CD3-NK\ Cells)}$ X 100 |
| CD8+ T regulatory cells | $\frac{(CD3+CD8+CD35+\ T\ Cells)}{(CD20+B\ Cells)+(CD3+T\ Cells)+(CD16+CD3-NK\ Cells)}$ X 100 |
| Classical monocytes (CM) | $\frac{(CD14+HLADR+\ CM)}{(CD14+HLADR+CM)+(CD16+HLADR+NCM)+(CD14+CD16+HLADR+\ IM)}$ X 100 |
| Non-Classical monocytes (NCM) | $\frac{(CD16+HLADR+\ NCM)}{(CD14+HLADR+CM)+(CD16+HLADR+NCM)+(CD14+CD16+HLADR+\ IM)}$ X 100 |
| Intermediate monocytes (IM) | $\frac{(CD14+CD16+HLADR+\ IM)}{(CD14+HLADR+CM)+(CD16+HLADR+NCM)+(CD14+CD16+HLADR+\ IM)}$ X 100 |
| CD20+ B Cell to CD3+ T Cell ratio | $\frac{(B\ Cells)}{(T\ Cells)}$ |
| CD4+ T cell to CD8+ T Cell ratio | $\frac{(CD4+T\ Cells)}{(CD8+T\ Cells)}$ |

**Supplementary Table 2.** Calculation of immune cell proportions or ratios.

| Linear models | | Sample size | Unique samples |
|---|---|---|---|
| **model 1** | *Principal component ~ age + sex + batch + 1\|animal ID* | 369 | 230 |
| **model 2** | *Principal component ~ age * sex + batch + 1\|animal ID* | | |
| **model 3** | *cell type/cell ratio ~ age + sex + batch + 1\| animal ID* | | |
| **model 4** | *cell type/cell ratio ~ age * sex + batch + 1\| animal ID* | | |
| **model 5** | *Principal component ~ status + age + sex + batch + 1\| animal ID* | 250 | 145 |
| **model 6** | *Principal component ~ status * age + sex + batch + 1\| animal ID* | | |
| | *Principal component ~ status * sex + age + batch + 1\| animal ID* | | |
| **model 7** | *cell type/cell ratio ~ status + age + sex + batch + 1\| animal ID* | | |
| **model 8** | *cell type/cell ratio ~ status * age + sex + batch + 1\| animal ID* | | |
| | *cell type/cell ratio ~ status * sex + age + batch + 1\| animal ID + 1\|Social Group* | | |
| **model 9** | *cell type/cell ratio in males or in females separately ~ status + age + batch + 1\| animal ID + 1\|Social Group* | Males 134 Females 116 | Males 73 Females 70 |

**Supplementary Table 3.** Linear mixed models used for data analysis using the *lmer* package in the R statistical software.

| **Cells** | Pearson's Correlation | **P value** |
| --- | --- | --- |
| B Cell/T Cell Ratio | -0.93 | $3.7 \times 10^{-160}$ |
| CD20+ B Cells | -0.86 | $1.3 \times 10^{-111}$ |
| CD4/CD8 T Cell Ratio | -0.5 | $4.4 \times 10^{-25}$ |
| Classical Monocytes | -0.28 | $3.0 \times 10^{-8}$ |
| NK Cells | -0.03 | 0.53 |
| CD4+ T Regulatory Cells | 0.24 | $3.2 \times 10^{-6}$ |
| Non-Classical Monocytes | 0.24 | $2.6 \times 10^{-6}$ |
| Intermediate Monocytes | 0.27 | $1.9 \times 10^{-7}$ |
| CD8+ T Regulatory Cells | 0.4 | $6.5 \times 10^{-16}$ |
| CD4+ T Cells | 0.5 | $4.3 \times 10^{-25}$ |
| CD8+ T Cells | 0.77 | $5.1 \times 10^{-74}$ |
| CD3+ T Cells | 0.86 | $1.3 \times 10^{-111}$ |

**Supplementary Table 4.** Cell type Pearson's correlation coefficient and p values with Principal Component 1.
